# MiR-195 regulates mitochondrial function by targeting mitofusin-2 in breast cancer cells

**DOI:** 10.1101/476812

**Authors:** Paresh Kumar Purohit, Ruairidh Edwards, Kostas Tokatlidis, Neeru Saini

## Abstract

Mitochondrial dynamics is a highly dysregulated process in cancer. Apoptosis and mitochondrial fission are two concurrent events wherein increased mitochondrial fragmentation serves as a hallmark of apoptosis. We have shown earlier that miR-195 exerts pro-apoptotic effects in breast cancer cells. Herein, we have demonstrated miR-195 as a modulator of mitochondrial dynamics and function. Imaging experiments upon miR-195 treatment have shown that mitochondria undergo extensive fission. We validated mitofusin2 as a potential target of miR-195. Which may provide a molecular explanation for the respiratory defects induced by miR-195 over-expression in breast cancer cells? Active, but not total, mitochondrial mass, was reduced with increasing levels of miR-195. We have further shown that miR-195 enhances mitochondrial SOD-2 expression but does not affect PINK1 levels in breast cancer cells. Collectively, we have revealed that miR-195 is a modulator of mitochondrial dynamics by targeting MFN2 thereby impairing mitochondrial function. Concomitantly, it enhances the scavenger of reactive oxygen species (SOD-2) to maintain moderate levels of oxidative stress. Our findings suggest a therapeutic potential of miR-195 in both ER-positive as well as ER-negative breast cancer cells.

## Introduction

Breast cancer is one of the leading causes of death in women worldwide. Treatment of breast cancer includes radiation, surgery and chemotherapy. The majority of breast tumors are hormone-dependent and they account for about 70% of breast cancers. About 80% of breast cancers are “ER-positive”, i.e. the cancer cells grow in response to the hormone estrogen ^1^. About 65% of these are also “PR-positive” (Progesterone receptor). Tumors that are *ER*/*PR-positive* are much more likely to respond to hormone therapy than tumors that are *ER*/*PR*-negative. Triple negative breast cancer (TNBC) lacks all three receptors i.e.ER, PR, and Her2. The Cancer therapy that targets these receptors does not work well with TNBC tumor ^2^. The development of drug resistance to chemotherapy is one of the major hurdles in treatment of breast cancer ^3^. The existing therapeutic approaches are not sufficient to root-out breast cancer. Consequently, development of novel drugs that target tumors irrespective of their hormone receptor status have a potential to strengthen the fight against breast cancer.

A growing number of studies have shown that microRNAs play key roles in the regulation of several cellular processes, whilst microRNA expression profiling has been associated with tumor development, progression and response to therapy ^4, 5^. MiRNAs are 20-25 nucleotide long sequences of non-coding RNA that regulate gene expression post transcriptionally. MiRNAs mimics and antagomiR have been widely used to normalize miRNA expression and hence gene expression networks in disease conditions ^6^. The ability of miRNA to restore gene expression to physiologically normal level renders microRNA-based strategies a potential therapeutic approach.

In several studies hsa-miR-195 has been reported to be differentially expressed in normal and breast cancer tissues ^7, 8, 9, 10^. A decrease in has-miR-195 expression in cancer tissue is due to increased methylation on CpG Island of miR-195 promoter ^11, 12^. Further, increased circulatory level of hsa-miR-195 in breast cancer patients and decline in circulatory miR-195 post-operation make hsa-miR-195 a potential biomarker for breast cancer ^13, 14, 15^. In previous studies we have unveiled a pro-apoptotic role of miR-195 in breast cancer and we have demonstrated that miR-195 downregulates Bcl2 by direct binding to its 3’UTR Sequence ^16^. In another study we have further shown the anti-proliferative, non-invasive and anti-metastatic role of hsa-miR-195 in breast cancer cells ^17^. These findings suggest hsa-miR-195 has strong anticancer properties with a promising therapeutic potential in breast cancer. Besides its pro-apoptotic role, hsa-miR-195 has also been shown to depolarize the mitochondrial inner membrane and to increase the calcium concentration in mitochondria ^17^. Despite all previous studies it remains unclear how miR-195 affects mitochondrial function. During our *in-silico* analysis we observed that mitofusin-2 (MFN2) acts as a predicted target of hsa-miR-195. Further reanalyzing our illumina microarray data, we found a differential expression of MFN2 upon over-expression of miR-195 in breast cancer cells. Herein, we have validated mitofusin-2 as a direct target of miR-195 in breast cancer cell lines. Over-expression of miR-195 in breast cancer cell induces mitochondrial fragmentation and diminishes the ability of mitochondrion to consume oxygen. The observed functional impairment of mitochondria induced by miR-195 is independent of the ER (Estrogen receptor) status of cells.

## Results

### MiR-195 down regulates MFN2 in Breast cancer cell lines

We have previously shown that miR-195 affects mitochondrial functioning by means of depolarization of the inner membrane and disturbing calcium homeostasis within the organelle ^17^. However the exact mechanism by which theses processes are controlled was not investigated. *In-silico* analysis using target scan revealed a miR-195 binding site in the 3’UTR of MFN2 (Figure1A). Mitofusin-2 is a crucial protein known to play a role in controlling mitochondrial dynamics and maintenance of mitochondrial calcium homeostasis. The miR-195 was upregulated using pSilencermiR-195 (miR) cloned vector whereas down regulated using antimiR-195 (AM)(Figure1B). As shown in Figure 1C, western blot analysis revealed significant down regulation (p-value<0.0005) of MFN2 upon over-expression of miR-195 in MCF-7 and MDA-MB-231 cells. While, MFN2 level was significantly increased (p-value<0.0005) when miR-195 expression was reduced in MCF-7 and MDA-MB-231 cell line. The 0.4 fold decrease (p-value<0.05) in luminescence was observed upon up-regulation of miR-195 in MDA-MB-231 cells whereas 0.16 fold decrease in luminescence was observed MCF-7 cells (Figure D) confirm direct binding of miR-195 to 3’UTR sequence of MFN2.

**Figure 1.**
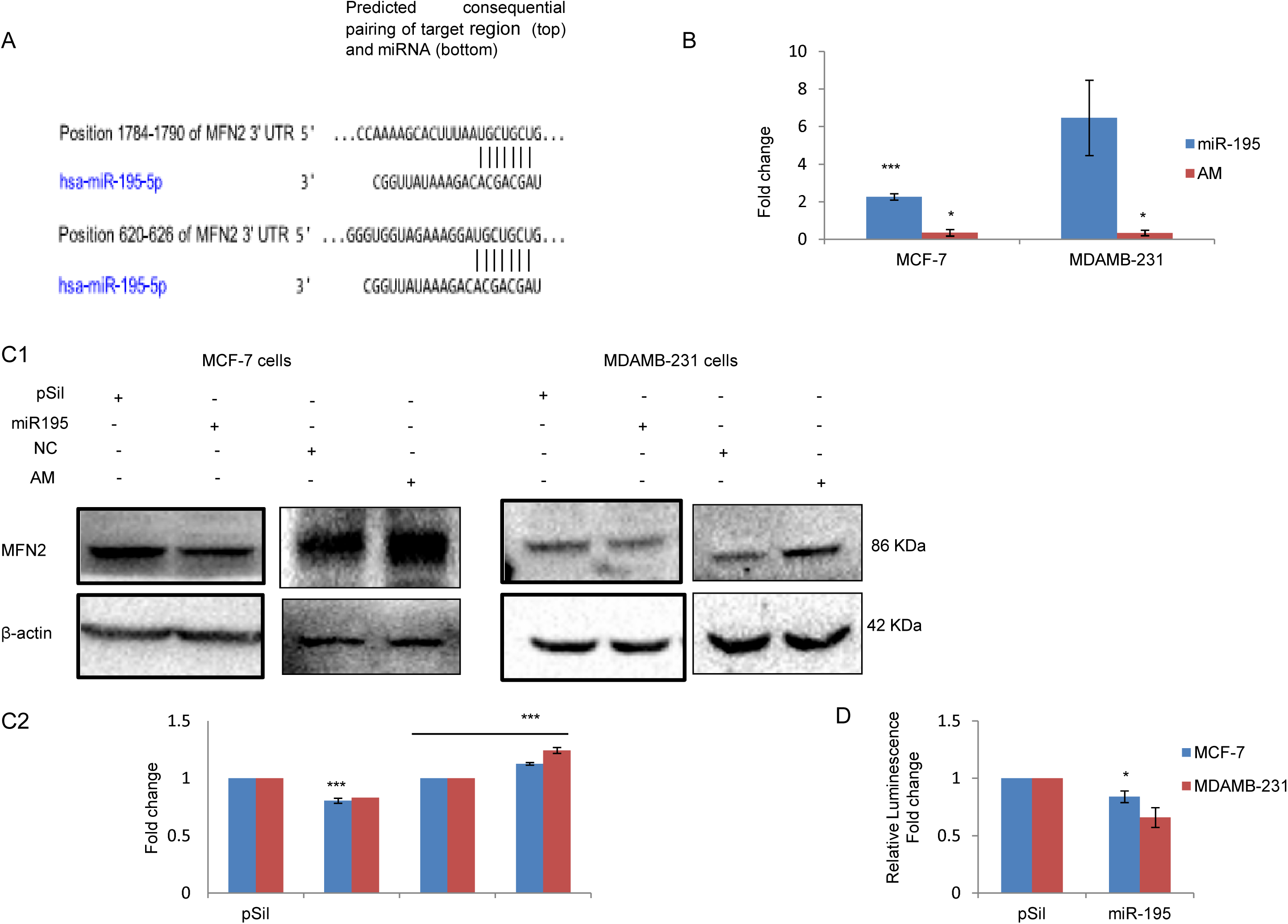
MiR-195 downregulates mitofusin-2 (mfn-2) by directly binding to its 3’UTR sequence. **(A)** Binding sites of miR-195 on 3’UTR sequence of MFN2 m-RNA **(B)** miR-195 over-expressed using p195 plasmid construct(miR-195) and knockdown using antimiR-195(AM) in MCF-7 and MDAMB231 cells **(C1)** miR-195 over-expression depleted MFN2 protein level while knockdown of miR-195 enhances MFN2 protein level in MCF-7 and MDA-MB-231cells **(C2)** mean fold change in MFN2 level ±SE for n=3 was plotted **(D)** Over-expression of miR-195diminshes relative luminescence in MDA-MB-231 cells, Representative plot of luciferase assay, mean fold change ±SE for n=3 was plotted, *: *p*<0.05, ***: *p*<0.001.

### MiR-195 affects mitochondrial morphology

Mitochondrial morphology is a critical factor that determines its activity. MFN2 has been shown to affect mitochondrial dynamics by promoting fusion events and hence mitochondrial morphology. Mitotracker CMXRos has been used to check morphology of the organelle. We observed increase fragmentation of mitochondria when we over-expressed miR-195 in MCF-7 and MDA-MB-231 cell lines (Figure 2). The highly networked tubular mitochondria in untreated cells tend to become rounded and small fragmented in shape upon over-expression of miR-195 in both cell lines, the complete fragmented morphology of mitochondria was observed in cells treated with CCCP i.e. the positive control (Figure 2).

**Figure 2.**
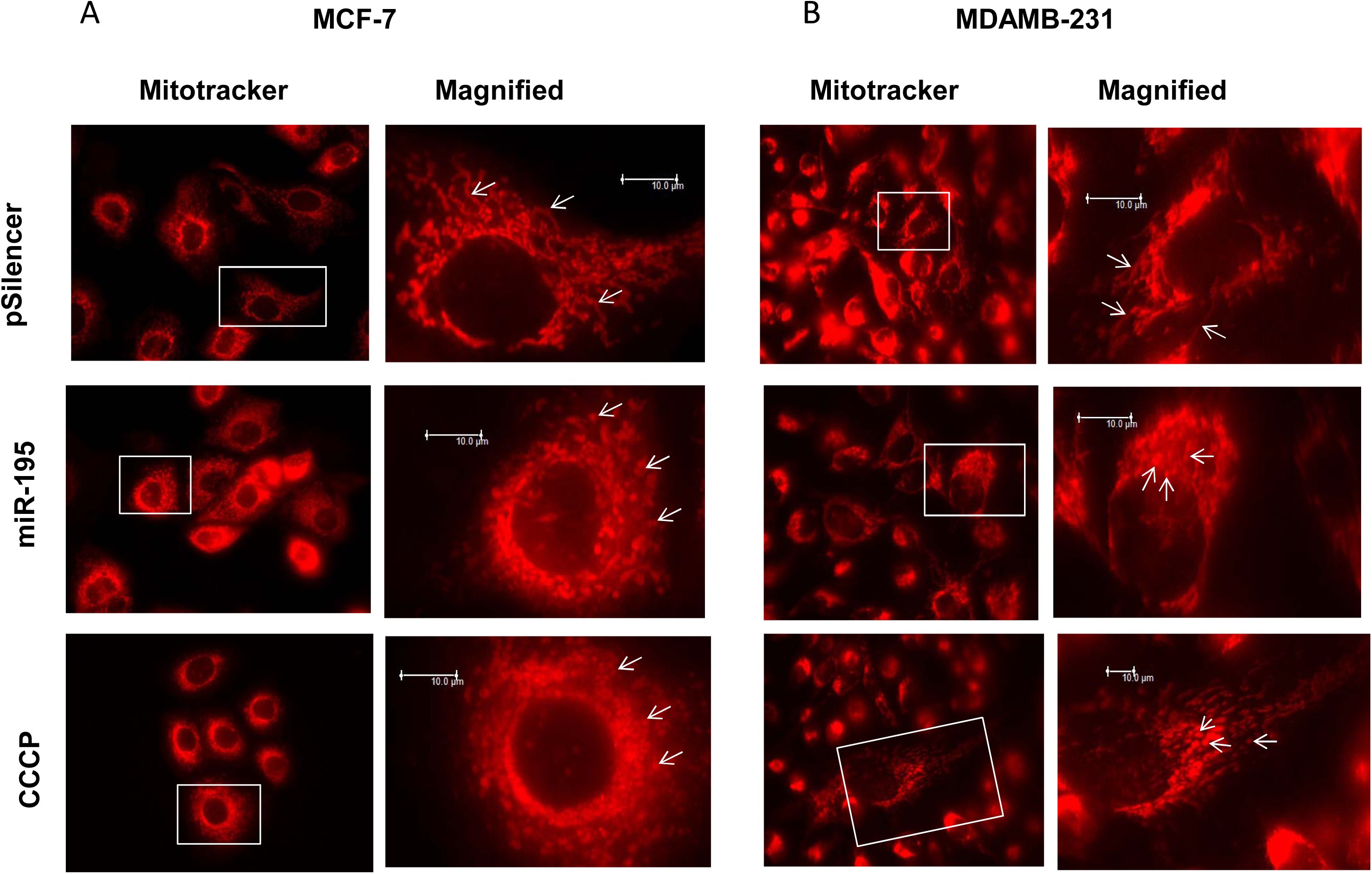
Mitofusin-2 down-regulation by miR-195 changes mitochondrial morphology and promotes mitochondrial fission. (A) Mitochondria become rounded and small fragmented upon up-regulation of miR-195 and gets severely fragmented upon CCCP treatment in MCF-7 cells **(B)** The highly elongated mitochondria becomes small tubular in shape upon up-regulation of miR-195and gets small fragmented upon CCCP treatment in MDA-MB-231 cells. Bar: 10µm.

### MiR-195 promotes mitochondrial fission by blocking fusion event

Mitochondrial morphology is critical for its functioning and it is regulated by mitochondrial dynamics. Apart from mitofusins there are other proteins whose levels change markedly during the mitochondrial fusion and fission events. Here we found that up-regulation of miR-195 affects the levels of both DRP1 (dynamin related protein) and OPA1. Specifically, in MCF-7 cells, DRP1 increases 1.9-fold (Figure 3A first panel) whilst OPA1 decreases 0.32-fold (Figure 3D first panel). In MDA-MB-231 cells 1.32-fold increase in DRP1 level was observed (Figure 3A second panel) whereas a 0.36-fold decrease in the OPA1 level (Figure 3D second panel)

**Figure 3.**
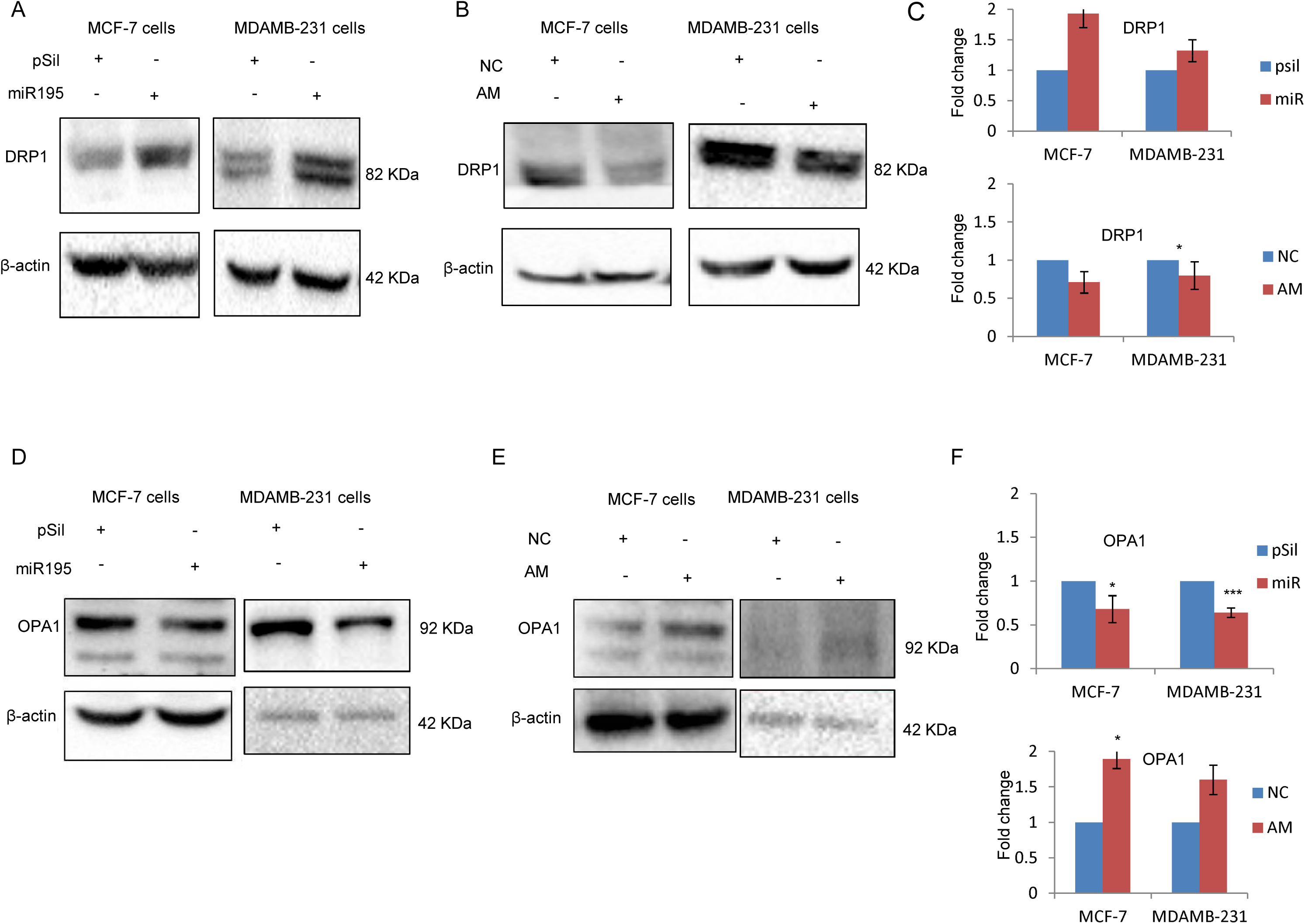
MiR-195 affects mitochondrial dynamics by blocking fusion. **(A)** Dynamin related protein (DRP1) gets upregulated upon over-expression of miR-195 in MCF-7 cells (right panel) and MDA-MB-231 cells (left panel) **(B)** Knockdown of miR-195 using antimiR-195(AM) reduces DRP1 protein level in MCF-7 cells (right panel) and MDA-MB-231 cells (left panel) **(D)** Mitochondrial inner membrane GTPase (OPA1) gets down-regulated upon over-expression of miR-195 in MCF-7 cells (right panel) and MDA-MB-231 cells (left panel) **(E)** Depletion of miR-195 using AM enhances OPA1 level in MCF-7 cells (right panel) and MDA-MB-231 cells (left panel) **(C, F)** mean fold change ±SE for n=3 was plotted, *: *p*<0.05, ***: *p*<0.001.

Conversely, knockdown of miR-195 has different effects. In MCF-7 cells DRP1 decreases 0.3-fold (p-value<0.05) (Figure 3B first panel) and OPA1 increases 1.9-fold (Figure 3E first panel). In MDA-MB-231 cells DRP1 decreases 0.2-fold (figure 3B second panel) whilst OPA1 increases 1.60-fold (Figure 3E second panel) The antimiR-195 treatment in both cell lines shows exactly opposite effects to that of miR-195. The enhanced DRP1 expression and reduced OPA1 protein level is a sign of miR-195-induced increase in mitochondrial fission events in breast cancer cell lines.

### MiR-195 diminishes active mitochondrial mass but does not promote degradation of mitochondria through mitophagy

Mitophagy and mitochondrial dynamics are two inter-connected process needed to maintain fully functional mitochondria by targeting defective and damaged mitochondria to a dedicated degradation process. Mitophagy is not the only process by which a healthy pool of mitochondria is maintained since damaged mitochondria can also undergo a repair pathway or they can fuse with completely healthy mitochondria and compensate the damage. As fusion events are blocked by the miR-195 by targeting MFN2 therefore fusion of damaged mitochondria with healthy ones is no longer an option for mitochondria to overcome the damaging oxidative stress. We have therefore checked for mitochondrial mass and mitophagy. The VDAC1 and mtHSP70 proteins are used as mitochondrial mass markers whilst PINK1 is used as a mitophagy marker. Upon over-expression of miR-195 or knockdown of miR-195 none of these markers (Figure 4A for PINK1, 4B for VDAC1, 4C mtHSP70 protein) have shown any significant difference compared to vehicle control in MCF-7 and MDA-MB-231 cell lines. These observations clearly suggest that miR-195 does not promote degradation through mitophagy of damaged mitochondria and hence total mitochondria mass does not change. We further probed for the functionally active mitochondria pool using mitotracker CMXRos (which is highly dependent on a functional mitochondrial transmembrane potential). We observed a decrease of active mass of mitochondria upon miR-195 treatment in MCF-7 and MDA-MB-231 cell lines. On the other hand, the mass of active mitochondria increased upon down-regulation of miR-195 in both cell lines. CCCP treatment was used as a positive control. Taken together, these observations suggest that miR-195-mediated mitochondrial membrane depolarization is not severe enough to induce mitophagy, but it is sufficient to diminish mitochondrial activity and hence the pool of active mitochondria.

**Figure 4.**
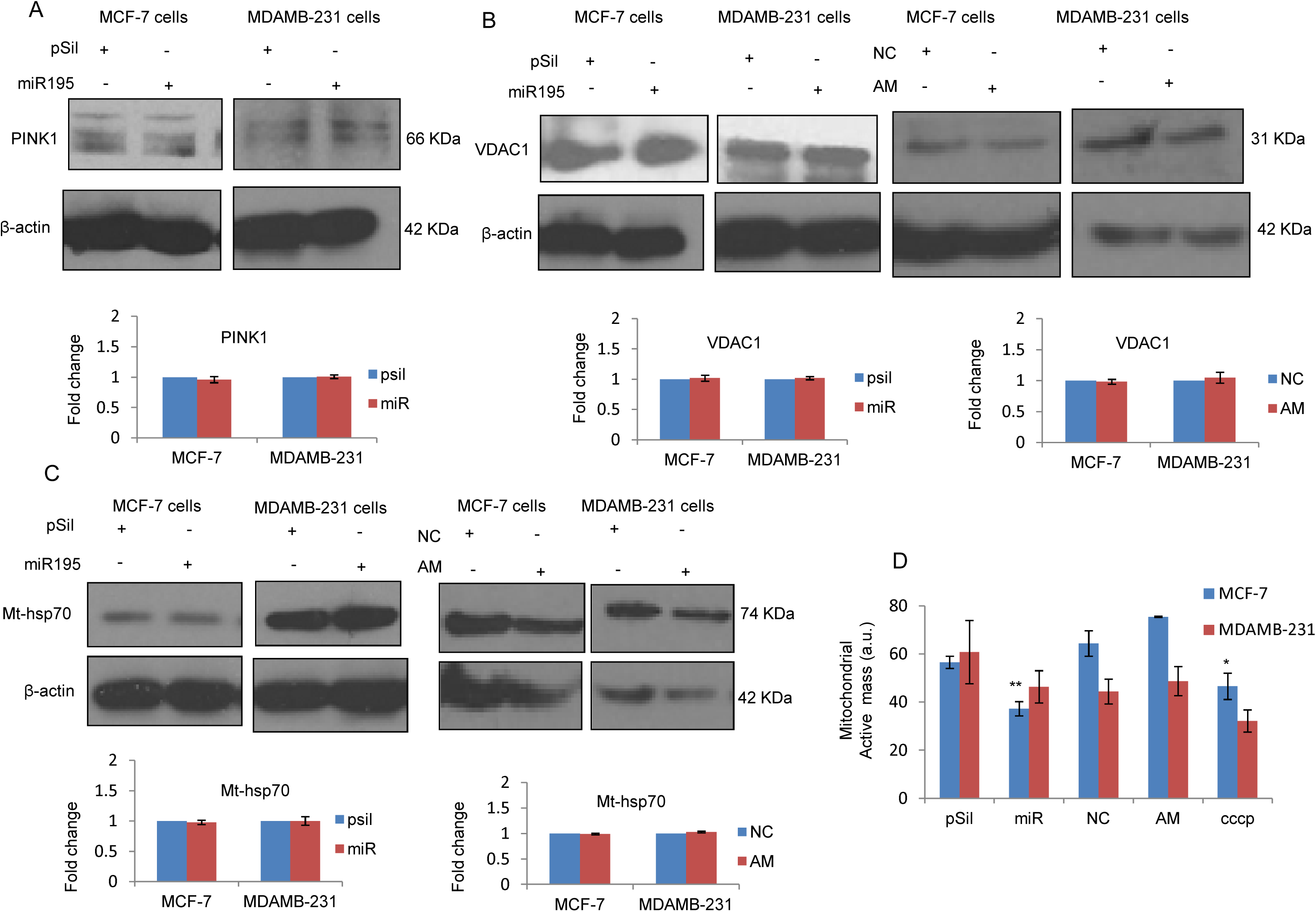
MiR-195 diminishes Active mitochondrial mass but not the total mass of mitochondria. **(A)** Mitophagy marker PINK1 level was not significantly changed with differential expression of miR-195 in MCF-7 (right panel) and MDA-MB-231 cells (left panel). The alteration in miR-195 level does not affect the protein level of **(B)** VDAC1 (mitophagy and mitochondrial mass marker) as well **(C)** mtHSP-70 (mitochondrial mass marker) in both breast cancer cell lines. Mean fold change ±SE for n=3 was plotted **(D)** Active mass of mitochondria was reduced upon miR-195 over-expression whereas active mass got resumed to normal level with depletion of miR-195 in MCF-7 and MDA-MB-231 cells, the positive control CCCP treated cells have shown lesser mitochondrial active mass, plot is representation of fold change in mean fluorescence intensity ±SE, *: *p*<0.05, ***: *p*<0.001.

### MiR-195 induces a defect in mitochondrial respiration

To further validate the effect of miR-195 on aerobic respiration, the Oxygen consumption rates of mitochondria (OCR) was measured directly with a SeaHorse-XF24 flux analyzer. The over-expression of miR-195 shows a potential respiratory defect as OCR is reduced, whilst down-regulation of mir-195 resulted in an increase in OCR in both the MCF-7 and the MDA-MB-231 cell lines. The OCR was used to further calculate other respiratory parameters such as basal respiration, maximal respiration, ATP production and spare respiratory capacity of mitochondria. miR-195 appears to attenuate basal respiration by 0.3 fold in MCF-7 and 0.35 fold in MDA-MB-231 cell lines whereas antimiR-195 treatment increases basal respiration by 1.24 fold and 1.27 fold in MCF-7 and MDA-MB-231 cell lines respectively (figure 5C), The maximal respiration also decreases by 0.35 fold and 0.3 fold in MCF-7 and MDA-MB-231 cell lines respectively, on contrary down-regulation of miR-195 enhances maximal respiration by 1.27 fold in MCF-7 and 1.47 fold in MDA-MB-231 cell lines (Figure 5D). As shown in figure 5E, miR-195 over-expression diminishes spare respiratory capacity by 0.33 fold and 0.51 fold in MCF-7 and MDA-MB-231 cell line respectively, on the other hand antimiR-195 increases spare capacity of mitochondria by 3.75 in MCF-7 and 2.16 fold in MDA-MB-231 cell lines. The OCR derived ATP production also decreases by 0.42 fold and 0.22 fold in MCF-7 and MDA-MB-231 cell lines respectively upon up-regulation of miR-195, conversely reduction in miR-195 level by antimiR-195 augments ATP production by 1.3 fold in MCF-7 and 3.72 fold in MDA-MB-231 cell line (figure 5F).

**Figure 5.**
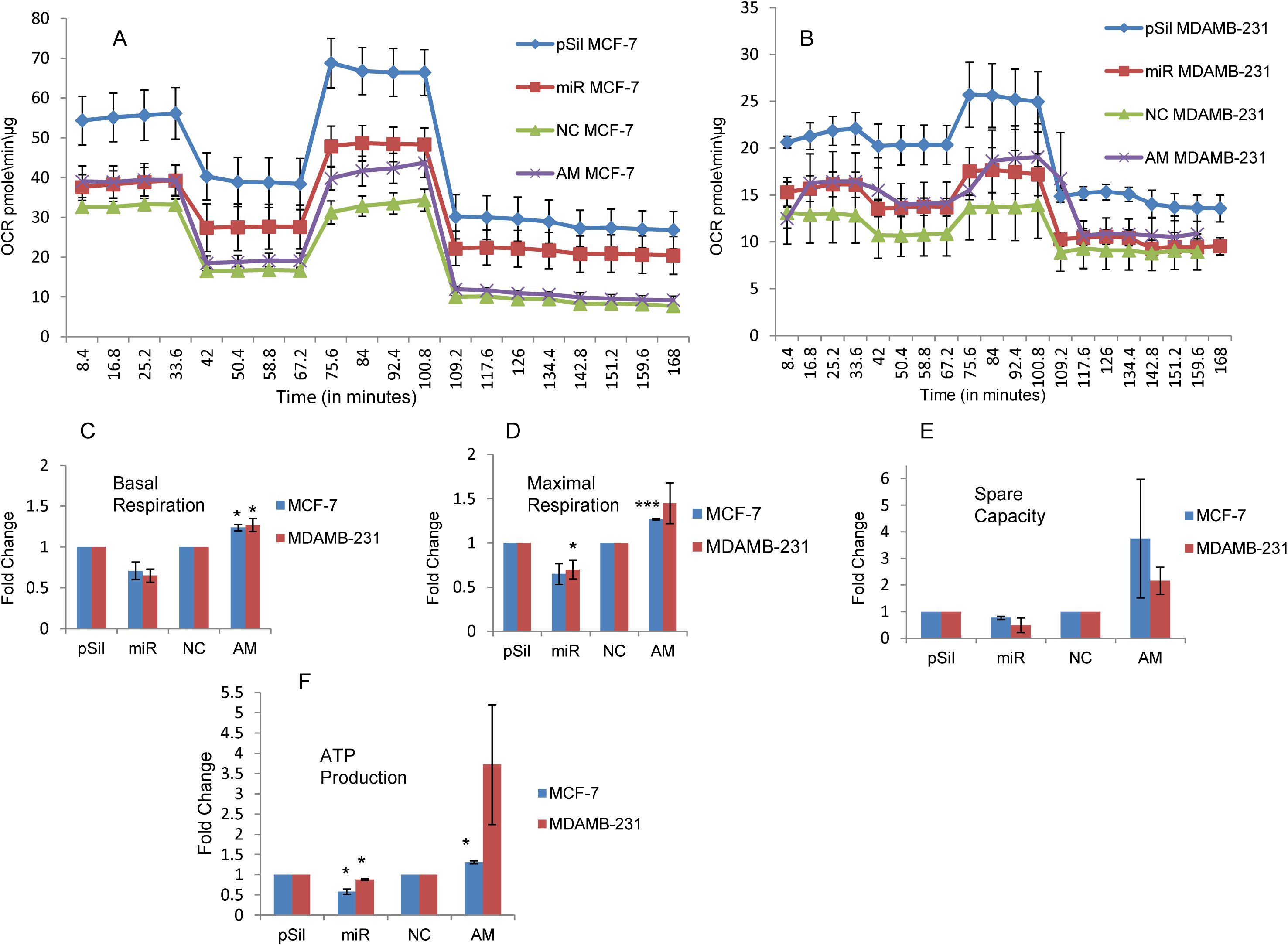
MiR-195 reduces OCR via MFN2 down-regulation. **(A)** Elevated level of miR-195 reduces OCR whereas declining miR-195 level augments OCR in **(A)** MCF-7 cells and **(B)** MDA-MB-231 cells, normalized OCR per min per microgram of protein ±SE from n=3 was plotted. The respiratory parameters such as **(C)** Basal respiration, **(D)** Maximal respiration, **(E)** Spare respiratory capacity of mitochondria and **(F)** ATP production was derived from OCR and mean fold change was plotted ±SE, *: *p*<0.05, ***: *p*<0.001.

### Oxidative stress is augmented by miR-195

The mitochondria are a prime source of energy production in cells, wherein, it utilizes the proton gradient across the membrane to generate ATP and convert molecular oxygen into water as a byproduct. The process of oxidative phosphorylation and electron transfer constantly takes place in inner mitochondrial membrane of metabolically active cells. The activity and proper assembly of electron transport chain complexes plays a central role in the electron flow and generation of the proton gradient across the membrane. Any defect in the ETC components leads to leakage of electrons to the mitochondrial matrix and that in turn converts molecular oxygen to superoxide, a damaging ROS. To check the effect of miR-195 on oxidative stress, we have probed for mitochondrial superoxide using Mitosox Red, a fluorescent probe that is rapidly oxidized by superoxides in mitochondria and produces red fluorescence. The over-expression of miR-195 enhanced superoxide levels 1.2-fold in MCF-7 cells and 1.13-fold in MDA-MB-231 cells. This increase in mitochondrial ROS suggests the involvement of ETC in the effect exerted by miR-195 on mitochondrial dynamics. To further examine the effects on ROS, we checked the expression of MnSOD2 as it is the key scavenger of superoxide in the matrix of mitochondria. Interestingly, the level of MnSOD2 increased 1.2-fold (p-value<0.05) in MCF-7 cells and 2-fold (p-value<0.01) in MDA-MB-231 upon over-expression of miR-195 (Figure 6A, 6B). Conversely, the MnSOD2 level decreased 0.3-fold (p-value< 0.05) in both MCF-7 and MDA-MB-231 cells when miR-195 was down-regulated (figure 6A, 6B). This additional evidence supports the role of miR-195 in maintaining the mitochondrial superoxide concentration at non-damaging levels by up-regulating its main scavenger protein (MnSOD2) and hence bypassing the mitophagy process.

**Figure 6.**
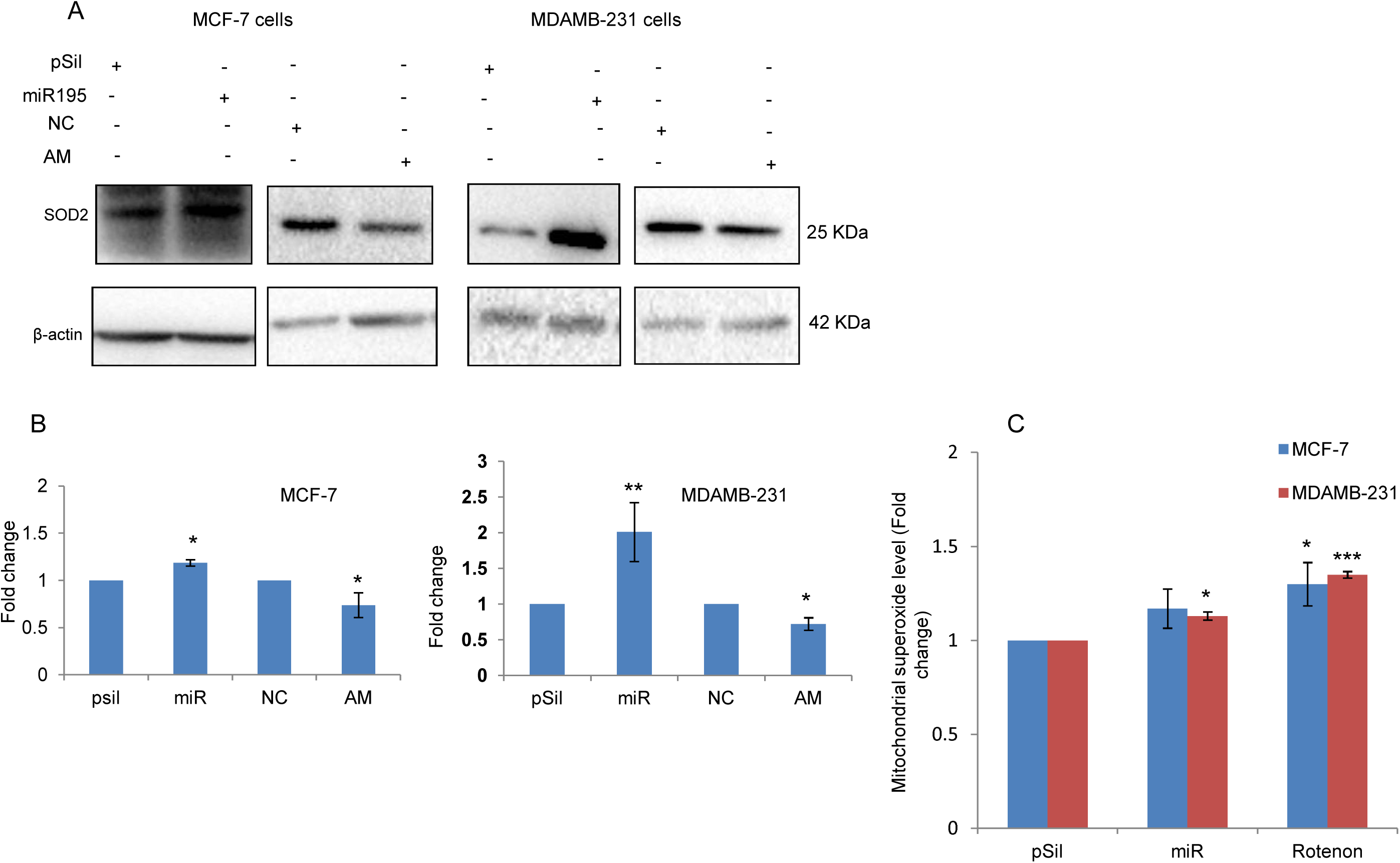
Moderate level of mitochondrial oxidative stress is maintained by miR-195. **(A)** Mitochondrial superoxide scavenger (mn-SOD2) elevates with miR-195 up-regulation whereas mn-SOD2 level was declined upon depletion of miR-195 in MCF-7 and MDA-MB-231 cells. (B) Mean fold change ±SE for n=3 was plotted. (C) mitochondrial superoxide level was augmented mildly in upon over-expression of miR-195 both breast cancer cells, relative fold change in mean fluorescence ±SE was plotted from n=3 for MDA-MB-231 and n=4 for MCF-7, *: *p*<0.05, ***: *p*<0.001.

**Figure 7.**
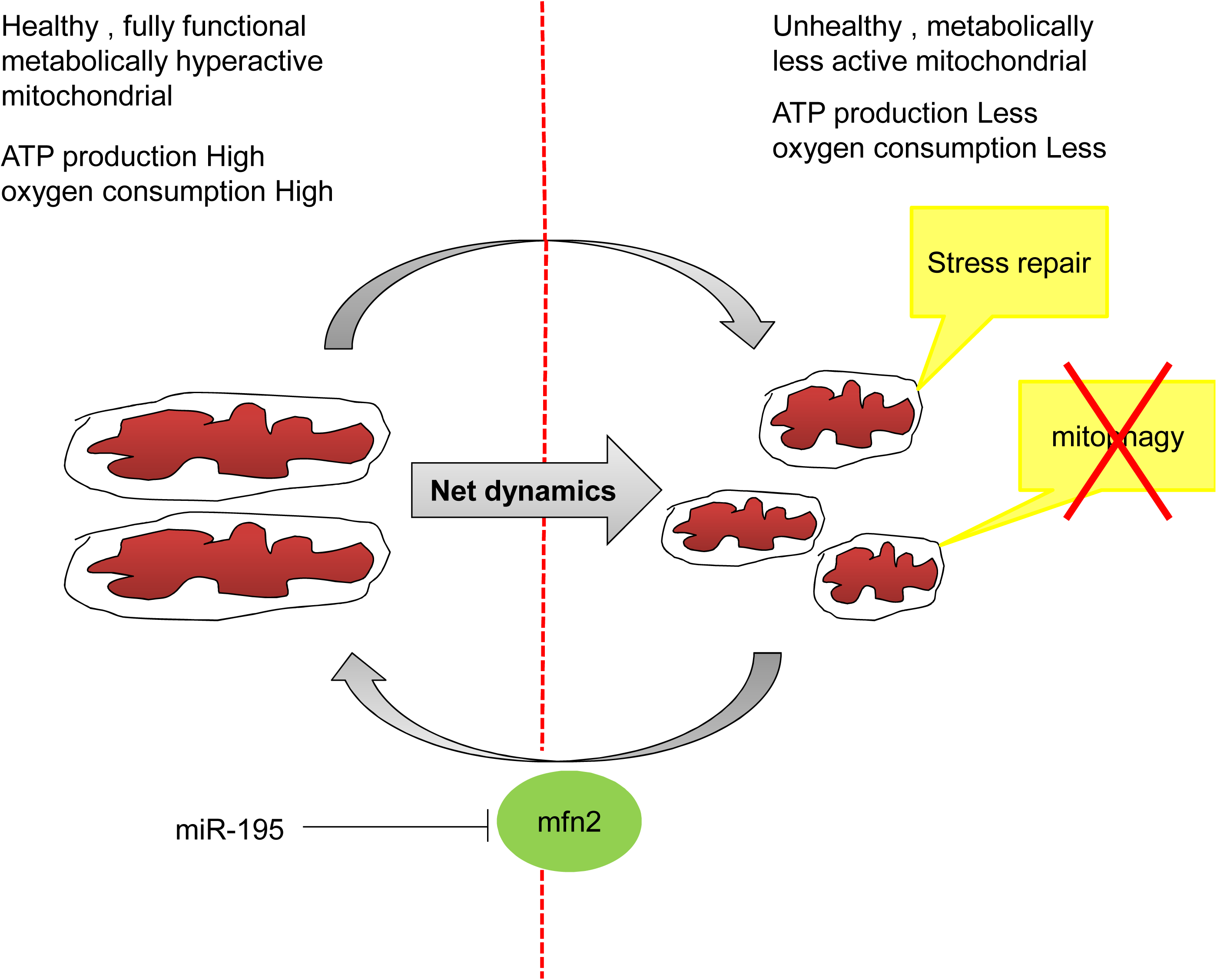
Schematic showing miR-195 mediated mitochondrial functional impairment via MFN2. MiR-195 affect mitochondrial dynamics by blocking MFN2 mediated mitochondrial fusion. The fragmented mitochondria as a result of miR-195 over-expression are metabolically less active with repairable oxidative stress.

## Discussion

Healthy mitochondria play a critical role in the determination of cell fate and the maintenance of cellular homeostasis. Mitochondrial dysfunction is a pathological hallmark for different diseases and stress-induced mitochondrial dysfunction is known to activate the intrinsic pathway of apoptosis ^18, 19, 20, 21^. The dynamic nature of mitochondria ensures their quality, whilst unhealthy mitochondria with irreparable damage undergo a specific degradation process called mitophagy ^22, 23, 24, 25, 26^. A healthy pool of mitochondria is maintained by a series of fusion-fission events. Mitofusin1 and Mitofusin2 facilitate fusion of the outer mitochondrial membrane ^27, 28, 29^, while the inner mitochondrial membrane gets fused by the inner membrane GTPase protein OPA1 ^30, 31, 32^. Mitochondrial fission is triggered upon localization of the DRP1 and FIS1 proteins on scissor site ^33, 34, 35, 36^. In conclusion, mitochondrial dynamics and mitochondria morphology are highly linked process that plays a major role in mitochondrial quality control to maintain cellular homeostasis.

Previously we demonstrated that hsa-miR-195 targets Bcl-2, it causes mitochondrial dysfunction, it induces apoptosis and augments the effect of etoposide in breast cancer cells. We further demonstrated that miR-195 also targets genes involved in de novo lipogenesis, it inhibits cell proliferation, migration, and invasion by targeting FASN, HMGCR, ACACA and CYP27B1 thereby potentially opening new avenues for the treatment of breast cancer. However, the underlying mechanism by which miR-195 causes mitochondrial dysfunction remains unexplored.

Herein, we first checked whether miR-195 can regulate mitochondrial shape and function. Our current findings revealed that miR-195 modulates mitochondrial shape and enhances fragmentation of mitochondria in breast cancer cells (Figure2). We also identified MFN2 to be a direct target of miR-195 in apoptotic breast cancer cell lines. The expression of MFN2 has been shown by other groups to be implicated in various cancers ^37^. The over-expression of MFN2 has been observed to enhance tumorigenesis while inhibition of MFN2 has been revealed to exert anti-cancer effects ^38^. Further, MFN2 has been shown to affect mitochondrial dynamics by promoting fusion events. In an independent study Leboucher and colleagues reported that the phosphorylation and proteosomal degradation of MFN2 under stress condition facilitates mitochondrial fission and apoptosis ^39^. Therefore targeting MFN2 using miR-195 expands the therapeutic potential of candidate microRNA.

We further demonstrated that miR-195 reduces the efficiency of mitochondria to consume oxygen. The observed differential levels of OPA1 and DRP1 upon alteration of miR-195 expression further suggested an indirect involvement of both proteins in modulation of mitochondrial dynamics by miR-195. Next, we were curious to know the fate of fragmented mitochondria in apoptotic breast cancer cells upon up-regulation of miR-195. Mitochondrial fission and increased oxidative stress is often known to decrease mitochondrial mass by committing damaged mitochondria to mitophagy. However, to our surprise, we did not observe any change in mitochondrial mass and hence no mitophagy but there was a decrease in the mass of active mitochondria. Taken together, these observations suggest that miR-195 affects mitochondrial functions and enhances oxidative stress but the stress is not sufficient to trigger mitophagy. It seems that miR-195 induces moderate oxidative stress to compromise mitochondrial activity and that is the reason behind a reduced mass of active mitochondria.

Further, to check how the moderate level of stress is being maintained by miR-195, we analyzed the mitochondrial MnSOD2 and observed significant up regulation of superoxide scavenging in mitochondria under the influence of miR-195. This illustrates a very subtle effect of miR-195 that enhances oxidative stress on the one hand by affecting mitochondrial dynamics, whilst it does not allow excess ROS accumulation by enhancing the level of the scavenger (MnSOD2).

The role of pro-apoptotic miR-195 in modulation of mitochondrial dynamics by targeting MFN2 renders miR-195 an effective anticancer molecule. Our results revealed miR-195 mediated potential respiratory defects in MCF-7 and MDA-MB-231 cell lines that characterize miR-195 as a modulator of mitochondrial respiration. The current study gives detailed mechanistic insight into the mitochondrial functional impairment by miR-195 in apoptotic breast cancer cell lines. The results presented here expand our previous finding that miR-195 mediates differential expression of components of the mitochondrial electron transfer chain in breast cancer cell lines ^17^. The present study provides a much needed framework to gain further insight into the role of miR-195 as a putative potential therapeutic molecule not just in cancer but various metabolic as well as neurodegenerative diseases where mitochondrial function is being compromised.

## Methods

### Cell culture

Human breast adeno carcinoma cell lines MCF-7 and MDA-MB-231 were procured from National Centre for Cell Sciences (NCCS), Pune, India and cultured using DMEM containing 10% (v/v) fetal calf serum, 100 Units/ml penicillin, 100 μg/ml streptomycin, 0.25 μg/ml amphotericin at 37 ° C in a humidified atmosphere at 5% CO2.

### MiR-195 over-expression and knockdown

Over-expression of miR-195 was achieved using p195 (237 bp sequence containing premiR-195 cloned in psilencer 4.1 vector purchased from Ambion, Austin, TX, USA) ^16^ and pSilencer 4.1 empty vector was used as a control. The miR-195 knockdown was done using AntimiR-195 oligos (AM17000) and AntimiR miRNA inhibitor negative control (AM17010). Both oligos were purchased from Ambion, Austin, TX, USA. Lipofectamine 2000 and Lipofectamine LTX-Plus (Invitrogen, CA, USA) was used for transfection according to manufacturer’s protocol. Cells were trypsinized and harvested 24 hours post transfection.

### Mitochondrial morphology

The mitochondrial shape was visualized using mitochondria specific probe mitotracker CMXRos ^40^. CMXRos is a fluorescent dye that is lipophilic and cationic in nature. CMXRos binds to the negatively charged matrix side of the mitochondrial inner membrane. Cells were stained with 200 nM CMXRos for 15 minutes at 37 ⁰C post miR-195 treatments in cell culture incubator. The mitochondrial proton gradient and ATP synthesis uncoupler Carbonyl cyanide *m*-chlorophenyl hydrazone (CCCP) was used at 50 µM concentration as a positive control to visualize complete fragmentation of mitochondria. The cells were visualized at 63X magnification using fluorescence microscope (Leica DMI6000, Germany)

### RNA isolation and TaqMan assay

The miRNA-195 overexpression and knockdown was analyzed using Taqman probes specific to miR-195. The taqMan miRNA assay probes (RT000494 and TM000494) were purchased from applied biosystems. The total RNA was isolated from MCF-7 and MDA-MB-231 cell lines 24 hours post transfections with p195, antimiR-195 and their respective controls. The Trizol reagent (Ambion by Invitrogen, CA, USA) was used as per manufacturer’s instructions to isolate total RNA. The Integrity of RNA samples were checked at 1% agarose gel in a TAE buffer system. Quantification was done using a nanodrop and absorbance measurements at 260 nm, 280 nm and 230 nm. Five hundred nanograms of RNA of each sample was used to synthesize c-DNA using RevertAid H minus first strand cDNA synthesis kit as per manufacturer’s protocol for 18s r-RNA, whereas miR195-specific cDNA was synthesized using miR-195 specific primer (RT000494) and the TaqMan reverse transcription kit (Applied Biosystems) as per manufacturer’s protocol. The expression of matured miR-195 was estimated using TaqMan miRNA assay probe (TM000494, Applied Biosystems), the 18s r-RNA (Ref-4333760F, Applied Biosystems) was used to normalize the miR-195 expression. Data analysis was done using Pfaffl’s method ^41^.

### Protein preparation and Western blotting

Cells were lysed with RIPA lysis buffer (50 mM Tris–HCl, pH 7.4, 150 mM NaCl, 1% NP40, 0.25% Na-deoxycholate, 1 mM EDTA) containing protease inhibitors (1 μg/ml aprotinin, 1 μg/ml pepstatin, 1 μg/ml leupeptin, 1 mM PMSF,1 mM sodium fluoride and 1 mM sodium orthovanadate) for 30 minutes on ice. The lysates were centrifuged at 16000 g for 30 minutes at 4 °C and the supernatant was collected. Protein concentration was estimated by the BCA (Sigma, USA) method. Equal amount of proteins (50-100 μg) were separated on 10%-12% sodium dodecyl sulphate polyacrylamide gel electrophoresis (SDS-PAGE). Separated protein was further transferred to PVDF membrane (Mdi; Advanced Microdevices, India). The membrane was blocked using 5% Bovine serum albumin (BSA) for 1 h at RT and incubated with the respective antibodies in 1% BSA for 16 hours at 4 °C, blot was washed three times using TBST, and then further incubated with the suitable secondary antibody for 1 hour at room temperature. DRP1, OPA1 and MFN2 antibody were purchased from Abcam (Cambridge, UK) and used at 1:1000 concentrations. VDAC1 antibody was purchased from cell signaling and used at 1:1000 concentrations. β-actin and GAPDH were obtained from Sigma (Sigma, USA) and used at 1:5000 concentrations. β–actin and GAPDH was used as protein loading control to normalize the results. The secondary antibodies were HRP-linked and used at 1:5000 concentration and blots were developed using enhanced chemi-luminiscence (Pierce, Amersham). Integrated density values were calculated using the Image-J software.

### Luciferase reporter assay

3′UTR of MFN2 gene was cloned in pSichek2 luciferase vector backbone (Promega, Madison, WI, USA) and 100 ng of plasmid was used along with miR-195 treatment in a 12-well cell culture plate. Luciferase activity was measured using the dual luciferase reporter assay system (Promega, Madison, WI, USA) as per manufacturer’s protocol.

### Oxygen consumption rate measurement

For metabolic flux measurement cells 60,000 to 80,000 cells were seeded in each well of a XF-24 plate, whilst for the plate that was used as a blank (A1, B4, C3, D6) no cells were seeded in the wells. Transfection of miR-195, AntimiR-195 and their respective controls was done using Lipofectamine 2000 (Invitrogen, CA, USA) reagent at 70% to 80% cell confluency. 24 hours post transfection the cells were washed with XF-24 assay media and incubated in the same buffer for 1 hour at 37 °C in a CO_2_-free environment. A pre-hydrated XF-24 cartridge was loaded with inhibitor in their respective chambers. A cartridge with utility plate was kept in XF-24 flux analyzer machine and the program was run. After the calibration step, the utility plate was replaced with the cell culture plate and the metabolic flux of the cells was measured. The protein estimation was done using the Bradford method post experiment to check the equal seeding density in each well of XF-24 flux analyzer. The data was normalized per microgram of protein^42^.

### Mitochondrial mass measurement

Cells were seeded in 12-well plates and treatment was given once the desired confluency was reached. 50 µM CCCP treatment for 2 hours was given as a positive control of mitochondrial mass reduction. Cells were washed post treatments, Staining was done using 200 nM mitotracker CMXRos for 15 min at 37 °C. Cells were washed using PBS and harvested. Cells were further acquired using FACS (EasyCyte Guava Technologies).

### Mitochondrial reactive oxygen species measurement

Mitochondrial superoxide was measured using MitosoxRed staining ^43^. The cells were seeded in 24 well plate and miR-195 treatment was given at 70% cell confluency. 1 µM rotenone was used as positive control to generate endogenous superoxide radicals. Mitochondrial superoxide was stained using 5 µM MitosoxRed for 20 minutes at 37°C. Cells were harvested using 0.5% trypsin and washed with PBS. The cells were acquired using FACS (EasyCyte Guava Technologies).

### Statistical Analysis

A two-tailed student t-Test was used for statistical analysis. Mean of a minimum of two or a maximum of four independent experiments ±SE is plotted. P-value less than 0.05, less than 0.01 and less than 0.001 is represented by *, ** and *** respectively on plots.

## Acknowledgement

We would like to thank Mauricio Cardenas-Rodriguez (KT laboratory) for help with some of the preliminary SeaHorse experiments and Prof Stephen Tait (Beatson Institute for Cancer Research – Glasgow) for a gift of MCF-7 and MDA-MB-231 cells.

## Funding

This study was supported by Genome dynamics in Cellular Organization, Differentiation and Enantiostasis (GENCODE) BSC0123 project from the Council of Scientific and Industrial Research (CSIR). Paresh Purohit is registered PhD student through Academy of Scientific & Innovative Research (AcSIR), India. PKP is thankful to CSIR for a research fellowship and for support through the Newton-Bhabha UK-India PhD studentship scheme (Award 269852058). Work in the KT laboratory was supported by the Scottish Universities Life Science Alliance and the Scottish Funding Council (grant HR07019), the Royal Society-Wolfson research merit award to K.T (grant WM120111) and the BBSRC (grant BB/R009031/1). RE is supported by a Lord Kelvin-Adam Smith PhD studentship from the University of Glasgow.

## Conflict of interest

The authors declare that they have no conflict of interest.

